# DNA damage-signaling, homologous recombination and genetic mutation induced by 5-azacytidine and DNA-protein crosslinks in *Escherichia coli*

**DOI:** 10.1101/2020.10.27.357855

**Authors:** Julie A. Klaric, David J. Glass, Eli L. Perr, Arianna D. Reuven, Mason J. Towne, Susan T. Lovett

## Abstract

Covalent linkage between DNA and proteins produces highly toxic lesions and can be caused by commonly used chemotherapeutic agents, by internal and external chemicals and by radiation. In this study, using *Escherichia coli*, we investigate the consequences of 5-azacytidine (5-azaC), which traps covalent complexes between itself and the Dcm cytosine methyltransferase protein. DNA protein crosslink-dependent effects can be ascertained by effects that arise in wild-type but not in *dcm*Δ strains. We find that 5-azaC induces the bacterial DNA damage response and stimulates homologous recombination, a component of which is Dcm-dependent. Template-switching at an imperfect inverted repeat (“quasipalindrome”, QP) is strongly enhanced by 5-azaC and this enhancement was entirely Dcm-dependent. The SOS response helps ameliorate the mutagenic effect of 5-azaC but unbalanced expression of the SOS-induced DNA polymerases, especially PolIV, stimulates QP-associated mutagenesis. In the absence of Lon protease, Dcm-dependent QP-mutagenesis is elevated, suggesting it may play a role in 5-azaC tolerance. Deletions at short tandem repeats, which occur likewise by a replication template-switch, are elevated, but only modestly, by 5-azaC. We see evidence for Dcm-dependent and-independent killing by 5-azaC in sensitive mutants, such as *recA*, *recB*, and *lon*; homologous recombination and deletion mutations are also stimulated in part by a Dcm-independent effect of 5-azaC. Whether this occurs by a different protein/DNA crosslink or by an alternative form of DNA damage is unknown.

**Highlights:**

- 5-azacytidine is broadly mutagenic and recombinogenic
- In E. coli, 5-azaC promotes genetic instability through Dcm methyltransferase.
- There are other, unknown lesions induced by 5-azaC besides Dcm/DNA crosslinks
- 5-azaC induces the SOS response, protecting cells from killing and genetic instability

## 1. INTRODUCTION^1^

DNA protein crosslinks (DPCs) are a common spontaneous source of DNA damage and can be formed by both nonenzymatic or enzymatic mechanisms. Nonenzymatic DPCs can be induced by radiation and a wide variety of chemicals (Barker et al. 2005). For example, reactive oxygen species and aldehydes, either encountered from the environment or endogenously formed by metabolic processes, can covalently link proteins to DNA. In addition, a number of conserved cellular enzymes are covalently linked to DNA as a reaction intermediate (Ide et al. 2011). Two examples are DNA topoisomerases and DNA cytosine methyltransferases (“CMeTs”). Certain drugs trap these normally short-lived covalent intermediates. The quinolone class of antibiotics and anticancer drugs etoposide and doxorubicin are topoisomerase type IIA poisons for bacterial and eukaryotic enzymes, respectively, and trap the covalent tyrosyl-toposiomerase/phosphoryl-DNA intermediate (also known as the “cleaved complex”). Camptothecin, another chemotherapeutic drug, is a similar poison for eukaryotic type IB topoisomerases (Drlica and Zhao 1997; Nitiss 2009; Pommier et al. 2010). In addition, cytosine methyl transferases (CMeTs) form a covalent complex with C6 of cytosine in their reaction to methylate the C5 of cytosine in DNA or RNA (Figure 1)(Santi et al. 1984; Friedman 1985). The nucleotide analog 5-azacytidine (Figure 1) inhibits the completion of the reaction, trapping the protein covalently attached to the nucleotide base.

**Figure 1.**
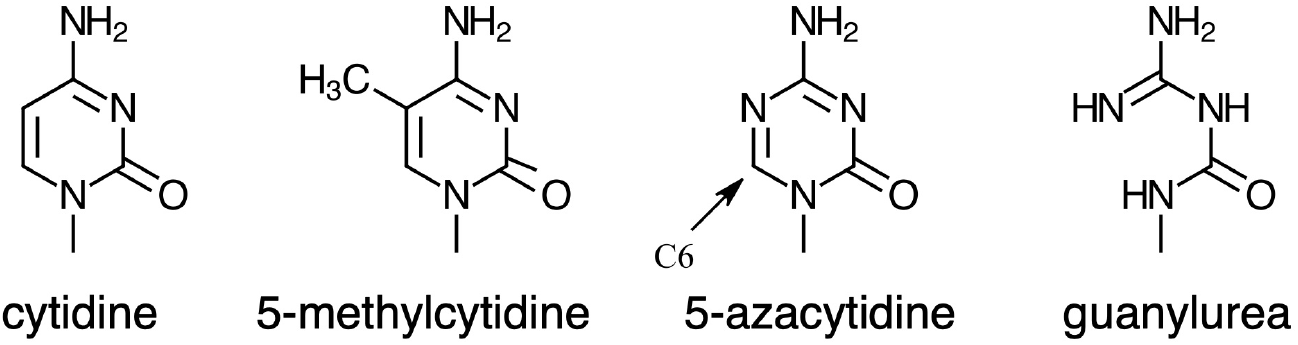
Cytidine-derived structures

DPCs impede both RNA transcription (Nakano et al. 2017) and replication fork progression (Kuo et al. 2007). In humans and other mammals, the inability to repair DPCs accelerates aging and leads to cancer proneness (Craft et al. 1987; Izzotti et al. 1999; Zahn et al. 1999; Wu et al. 2002; Swenberg et al. 2011; Garaycoechea et al. 2012).

In bacteria, DPCs caused by topoisomerase II poisons or 5-azacytidine can lead to double-strand breaks (DSBs). Accordingly, mutants in double-strand break (DSB) repair factors RecA and RecB are sensitive to both to quinolones and 5-azaC (Bhagwat and Roberts 1987; Lal et al. 1988; Butala et al. 2009; Salem et al. 2009; Ide et al. 2011), although there may be other toxic lesions in additional to DSBs. For topoisomerase II poisons, a cytotoxic lesion is some processed form of the trapped “cleaved complex” since killing can be suppressed by inhibitors of RNA and protein synthesis (reviewed nicely in (Drlica and Zhao 1997)). Even in the absence of frank DSB formation, these covalent complexes can block the completion of replication, allowing a second replication fork converging onto the site to produce a broken chromosome (Cox et al. 2000).

In bacteria, 5-azaC is a useful tool to study effects of DPCs in vivo whose targets are C5-cytosine methyl transfereases (CMeT) in restriction/modification systems. Restriction/modification systems are widespread throughout bacteria, and consist of an endonuclease component and a methyltransferase (Roberts et al. 2015). These systems are considered to be primitive innate immune systems, leading to the destruction of invading foreign DNA. A proportion of restriction systems use C5-cytosine methylation (Fig. 1) to provide immunity to restriction endonuclease cleavage (other systems methylate the N6 of adenine or the N4 of cytosine to confer immunity). Because many of these modification enzymes methylate specific DNA recognition sequences and are nonessential in the laboratory environment, we can target DPCs to particular sites to study their consequence in vivo. In addition, in the absence of the methyltransferase the adduct will not form, providing confidence that any effect is due to the DPC alone. We have recently documented that DPCs in *Escherichia coli* dramatically stimulate a particular class of mutation. This class of mutation arises by a replication template-switch reaction in an imperfect inverted repeat sequence (or “quasipalindrome”, “QP”)(Ripley 1982; Glickman and Ripley 1984; Yoshiyama and Maki 2003; Dutra and Lovett 2006; Lovett 2017). QPs often constitute mutational hotspots, both in bacteria and in eukaryotes (Hampsey et al. 1988; Greenblatt et al. 1996; Bissler 1998; Viswanathan et al. 2000; Yoshiyama et al. 2001; Yoshiyama and Maki 2003). Using QP mutational reporter strains, we discovered that the DPC-producing agents formaldehyde, ciprofloxacin and 5-azaC are potent mutagens for this class of mutations (Laranjo et al. 2018). For 5-azaC, the mutagenic effect requires *E. coli*’s endogenous CMeT, Dcm, indicating that it is the trapped DPC that induces mutation at the QP site (Laranjo et al. 2018).

Surprisingly, although the genes required in DSB repair, *recA* and *recB*, are necessary to avoid to the lethal effects of DPCs, they are not required for their mutagenicity (Laranjo et al. 2018). This suggests that the mutagenic lesions include something other than DSBs. That the mutagenic lesion may be a block to the DNA replication fork is suggested by the fact that impairment in the DnaB fork helicase greatly elevates QP-associated mutation (QPM)(Laranjo et al. 2018). To further explore how DPCs lead to genomic instability, we examine here the consequence of DPCs on genetic recombination and DNA damaging signaling. We further explore the effect of the SOS response, translesion DNA polymerase and the Lon and ClpP proteases on mutagenicity of 5-azaC in *E. coli* at a QP site. We additionally quantify effects on genomic rearrangements at short tandem repeats, which are caused by a different type of template-switching during DNA replication (Lovett et al. 1993; Lovett and Feschenko 1996).

This work finds that DPCs are broadly mutagenic, recombinogenic and inducers of the SOS response. The SOS response is protective in the avoidance of QPM, although unbalanced expression of translesion DNA polymerases, particularly DNA polymerase IV (DinB), is mutagenic. Most of the effects of 5-azaC are due to DPCs formed by the *E. coli* CMeT, Dcm; however, we also find evidence for Dcm-independent DNA lesions that are lethal or mutagenic in certain sensitive genetic backgrounds.

## 2. MATERIALS AND METHODS

### 2.1 Strains and growth conditions

All of the strains used are derivatives of *Escherichia coli* K-12 MG1655 (Bachmann 1972; Table 1). Isogenic strains of the indicated genotypes were constructed by P1vir transduction. DNA transformation by electroporation (Dower et al. 1988) was used to introduce plasmids and oligonucleotides for recombineering. The strains were grown at 37 °C in Luria broth (LB, Lennox formulation) medium, consisting of 1% Bacto-tryptone, 0.5% yeast extract, 0.5% sodium chloride and, for plates, 1.5% Bacto-agar. Tetracycline (Tc, 15 μg/mL), chloramphenicol (Cm, 30 μg/mL), ampicillin (Ap, 100 μg/mL) and kanamycin (Km, 45-60 μg/mL) were used for genetic selections. Lac^+^ reversion mutants were selected on lactose minimal medium (Lac Min XI) containing 56/2 salts (Willetts et al. 1969), 0.2% lactose (Sigma Aldrich, St. Louis, MO, USA), 0.001% thiamine (Sigma Aldrich), and 2% agar (ThermoFisher, Sparks, MD, USA). X-gal (40 μg/mL) and IPTG (0.1 mM) (both from Gold Bio, St. Louis, MO, USA) were included in lactose selection medium as a visual aid for counting colonies. Glucose minimal (Glu Min XI) control plates were made similarly but contained 0.2% glucose in place of lactose.

**TABLE 1:**
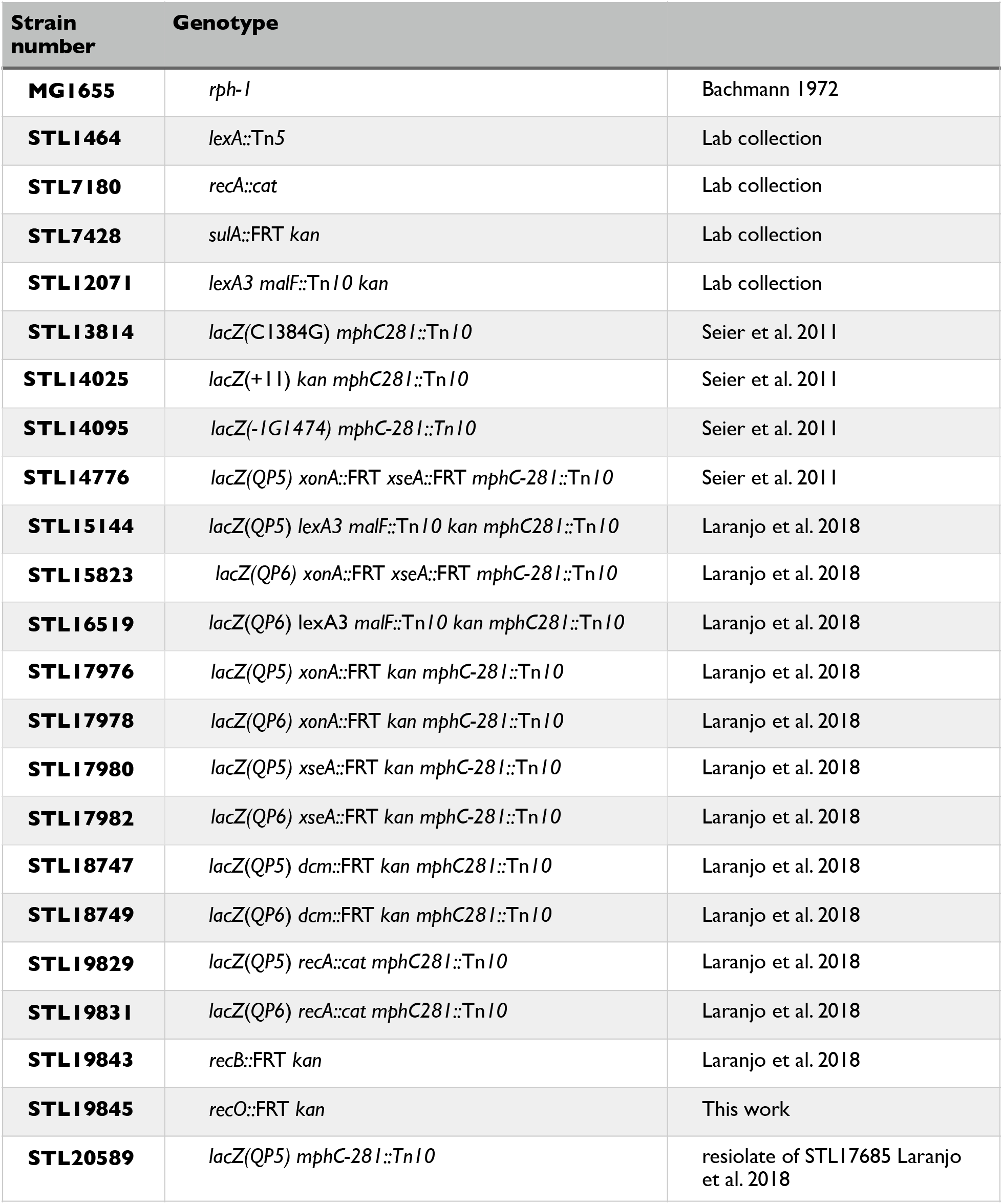
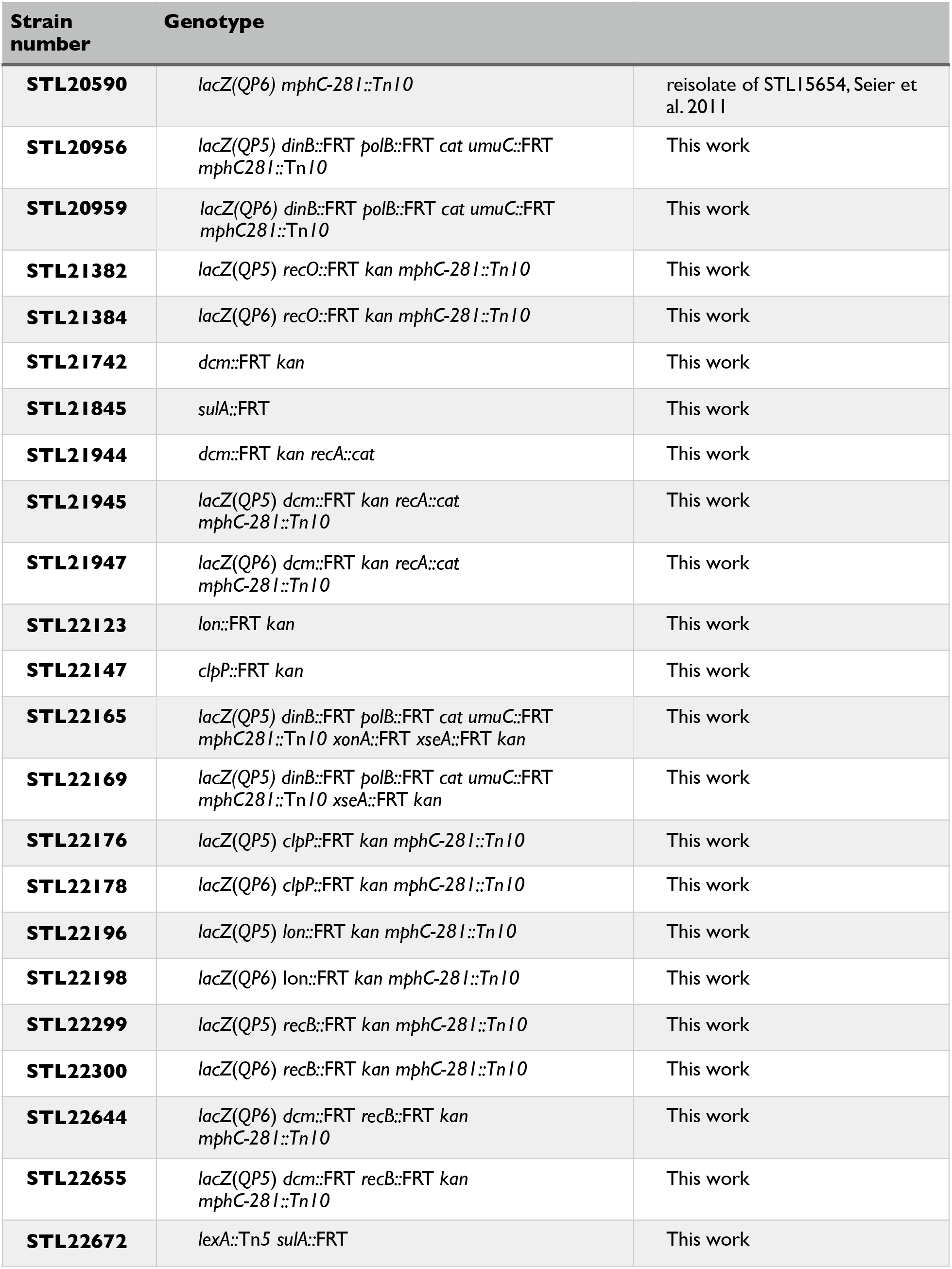
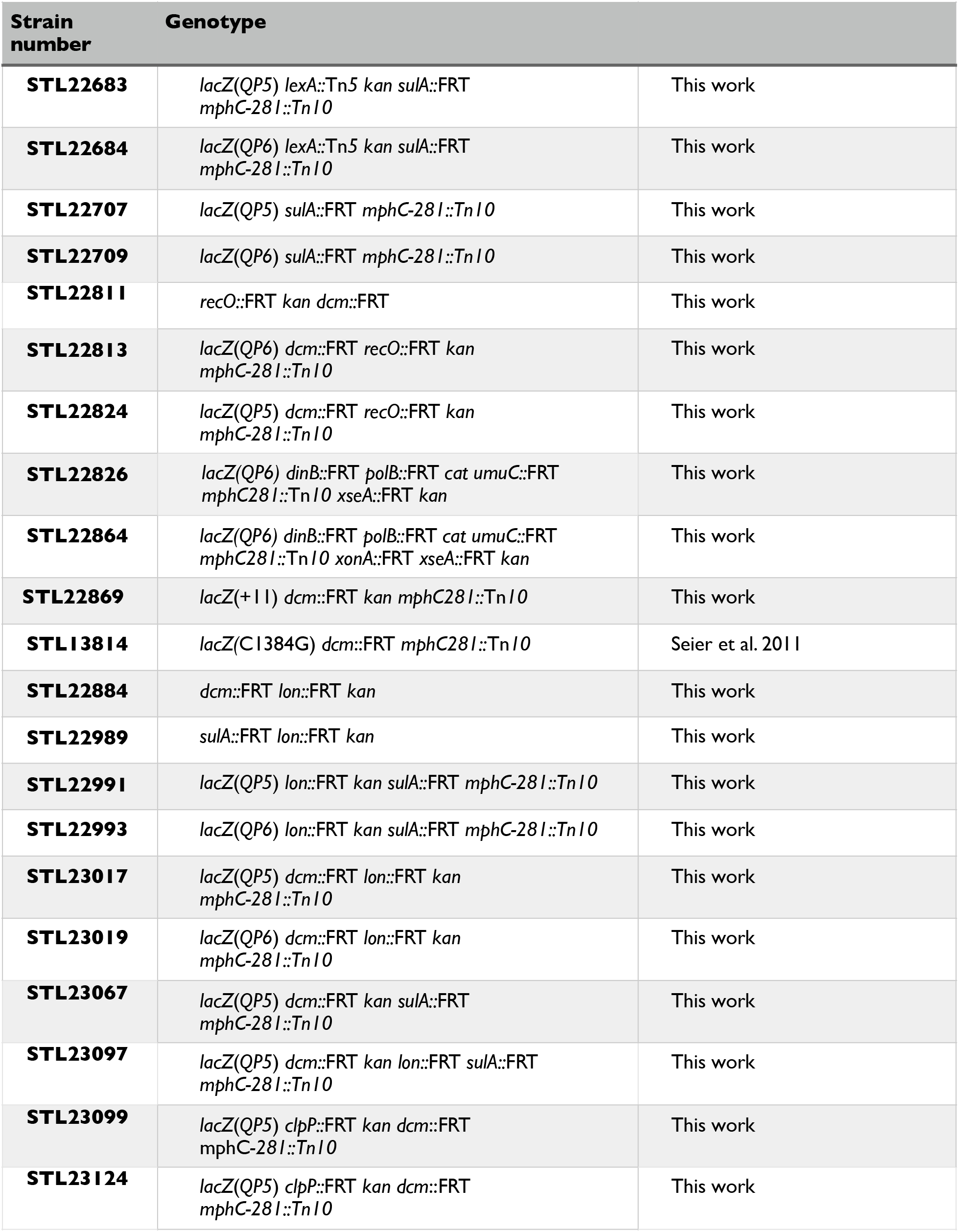
*E. coli* K-12 strains used in this study

### 2.2 Survival assays

For survival to 5-azacytidine (5-azaC, Sigma Aldrich), MG1655 and isogenic derivative (Table 1) were grown to mid-log phase (OD6_00_ of approximately 0.2-0.4), then split and treated at the indicated concentrations or untreated for 2 hr prior to serial dilution and plating on LB media. Survival experiments were performed for at least 3 replicates.

### 2.3 Luciferase assays

To measure induction of the SOS response, we used luciferase fusions to the *recA* and *dinB* promoters as previously described (Goldfless et al. 2006) and strain backgrounds MG1655, STL19845, STL21742, and STL22811. A single colony was used to inoculate 2 mL of LB+Ap media and grown at 37° with agitation for 2 hours until OD6_00_ ≈ 0.5. The culture was then diluted 1:100 in fresh LB+Ap media and grown again at 37C with agitation for 2 hours. The culture was then diluted 1:100 into fresh LB+Ap media in a Costar 96 Well Assay Plate (Treated Polystyrene, Black Plate, Clear Bottom). The plate was then grown in a BioTek Cytation 1 Plate Reader at 37° shaking for 75 minutes, with OD6_00_ and luminescence readings being taken every 15 minutes. After 75 minutes, 5-azacytidine was added to the appropriate wells to a final concentration of 12.5 ug/mL. Cultures were then grown for 3.5 hours shaking at 37° with OD6_00_ and luminescence readings taken every 15 minutes. Relative luminescence units (RLU) values were calculated by normalizing the luminescence readings (in counts per second) to the OD6_00_ of the cultures.

### 2.4 Deletion assays

Deletion of 101 bp tandem repeat in *tetA* was performed as described using plasmid pSTL57 (Lovett et al. 1994) and strains MG1655 and STL21742. Deletion of 11 bp tandem repeats in *lacZ* was performed as described (Seier et al. 2011) with strain STL14025 For experiments involving 5-azacytidine (Sigma Aldrich) or formaldehyde (ThermoFisher), mid-log phase cultures were split, and treated with the drug, or not, for 2 hours at the indicated concentrations before serial dilution and plating on appropriate selective media. Data are presented as the fold-change in deletion frequencies of the treated vs. untreated samples.

### 2.5 Crossover recombination assay

Recombination between 411 bp of homology in plasmids pSTL330 and pSTL36 was selected by Tc-resistance as described (Lovett et al. 2002) and strain backgrounds MG1655, STL21742 and STL19845. Mid-log phase cultures were split, and for two hours were treated with 5-azaC at the indicated concentrations before serial dilution and plating on appropriate selective media. Both wt and *recO* mutants were assayed with 4 independent replicates.

### 2.6 Gene conversion recombination assay

Gene conversion is selected in this assay by the Lac^+^ progeny of a strain carrying two mutant alleles of *lacZ*, one with an internal deletion at the natural *lacZ* locus (7.83 centisomes) and an internal 500 bp *lacZ* fragment integrated at *att*Tn7 near *glmS* (84.28 centisomes). The *lacZ* recipient allele was constructed by recombineering using a single-strand DNA oligonucleotide and DNA transformation (Sharan et al. 2009). The 5 nucleotides spanning the active site amino acid codon, from nucleotides 1384-1388 of the *lacZ* ORF, were deleted by transformation of oligonucleotide 5’ ATCACCCGAG TGTGATCATC TGGTCGCTGGG GAATAGGCCA CGGCGCTAAT CACGACGCGC TGTATCGC in a MG1655 strain carrying the pSIM6 recombineering plasmid (Sharan et al. 2009), yielding strain STL20478 after curing the pSIM6 plasmid with growth at 42°. To construct the recombination donor locus, we used a Tn7-based based system to introduce a DNA fragment carrying 250 bp of homology flanking both sides of the *lacZ* active site deletion into the *att*Tn7 site, near *glmS* (McKenzie and Craig 2006). The DNA fragment produced by colony PCR with oligonucleotides 5’ GGGGcccggg GCTGATGAAG CAGAACAACT TTAACGCCGT 3’ and 5’ GCttaattaa ACTGTTACCC ATCGCGTGGG CGTATTCGCA 3’ was inserted into temperature-sensitive Tn7 vector pGRG25 (McKenzie and Craig 2006) after cleavage with PacI and XmaI (New England Biolabs). Transformation of this plasmid into strain STL18091 (MG1655 *bglG*::FRT *kan*) was selected by Ap-resistance at 30°, after which the plasmid was cured by growth in LB at 42°; integrants of the Tn*7* end-flanked fragments were identified by PCR among survivors as described (Laranjo et al. 2017). This allele was transduced into strain STL20478 by cotransduction with *bglG*::FRT *kan*. The *kan* was then removed using FLP plasmid pCP20 (Datsenko and Wanner 2000). The resulting recombination assay strain carrying both loci (attTn7::’*lacZ*’ *lacZ*_*as*Δ_) is STL22043. Homologous recombination in this strain generates a *lacZ^+^* locus in its natural site; the donor locus at *att*Tn7 is unchanged. Strains carrying only the donor or only the recipient *lacZ* locus never yielded Lac^+^ progeny, nor did *recA* strains carrying both alleles, confirming that *lacZ^+^* isolates arise by homologous recombination between the loci.

For the recombination assays, cultures were grown in LB to midlog phase, then concentrated 5x by centrifugation and resuspension into LB medium containing 5-azaC or not. Growth of these split cultures was continued for 2 hours after which they were harvested by centrifugation, washed with 1 x 56/2 salts and serially diluted. Dilutions were plated on Lac Min XI to determine the number of Lac^+^ recombinants and on Glucose Min XI and LB for total number of CFU. LB plates were incubated at 37° 1 day and minimal plates for 3-4 days.

### 2.7 Mutation assays

We used a chromosomal *lacZ* mutational reporters specific for G to C transversion and for quasipalindrome-associated template-switch mutations, QP5 and QP6, previously described (Seier et al. 2011). The strains used are listed in Table 1, with the genotype including *lacZ*(QP5) or *lacZ*(QP6). C to G transversion mutations were measured using STL13814 ((Seier et al. 2011) and a *dcm*::FRT *kan* transductant of this strain, STL22870. +1 G frameshift mutations were measured using strain STL14095 carrying the indicated plasmids. 5-azaC effects were determined with mid-log phase split cultures and with treatment for 2 hours prior to dilution and plating.

### 2.8 SOS polymerase plasmids

GATEWAY (Thermo Fisher) recombinational cloning technology was used to clone *dinB, polB* and *umuCD’* into the pBAD18 arabinose-controlled expression vector. From MG 1655 colonies, the *dinB* gene was amplified with the following PCR primers 5’ GGGGACAAGT TTGTACAAAA AAGCAGGCTT CGAAGGAGAT AGAACCatgc gtaaaatcattca and 5’ GGGGACCACT TTGTACAAGA AAGCTGGGTC tcataatccc agcacc, the *polB* gene with 5’ GGGGACAAGT TTGTACAAAA AAGCAGGCTT CGAAGGAGAT AGAACCgtgg cgcaggcagg ttttatc, 5’ ggggACCACT TTGTACAAGA AAGCTGGGTC tcaaaatagc ccaagttgccc. The *umuD’C* gene was amplified from plasmid MP1 (Addgene) with primers 5’ GGGGACAAGT TTGTACAAAA AAGCAGGCTT CGAAGGAGAT AGAACCatga agagattgca gctcatg, 5’ ggggACCACT TTGTACAAGA AAGCTGGGT Cttatttgac cctcagtaaa tcag and inserted using BP reactions into pDONR201. LR reactions transferred these insert to the pSTL360 plasmid ‘(Dutra and Lovett 2006), a destination plasmid derived from pBAD18 (Guzman et al. 1995).

## 3. RESULTS

### 3.1 Recombination pathways and DPC processing

The *Escherichia coli* K-12 genome encodes one cytosine methyl transferase, Dcm (“<u>D</u>NA <u>c</u>ytosine <u>m</u>ethylase”). The function of this gene is not known but it is likely a remnant of a restriction/modification system. Dcm methylates the second C in the sequence “CC[A/T]GG”; this is the same specificity as the EcoRII restriction/modification system, encoded on an *E. coli* antibiotic-resistance conferring-plasmid, derived from a clinical isolate (Bannister and Glover 1968; Yoshimori et al. 1972; Bhagwat et al. 1986; Bhagwat et al. 1990).

*E. coli* K-12 wild-type cells are not particularly sensitive to killing by 5-azaC and mutants in *dcm* are only modestly more resistant (Fig. 2). The fraction of potential sites methylated by Dcm is unknown but there is evidence that not every potential Dcm site is methylated (Bhagwat et al. 1990; Ringquist and Smith 1992); for example, EcoRII overexpression confers additional sensitivity to 5-azaC in *dcm+ E. coli* (Bhagwat et al. 1990; Krasich et al. 2015). With only the endogenous Dcm CMeT, *recA* and *recB* mutants, deficient in DSB repair, show enhanced sensitivity (Fig. 2), confirming previous studies (Bhagwat and Roberts 1987; Lal et al. 1988; Butala et al. 2009; Salem et al. 2009; Ide et al. 2011). This sensitivity is partially dependent on Dcm. There is, however, residual sensitivity, suggesting Dcm-independent toxicity of 5-azaC. This residual sensitivity of *recA dcm* has also been previously documented (Bhagwat and Roberts 1987).

**Figure 2.**
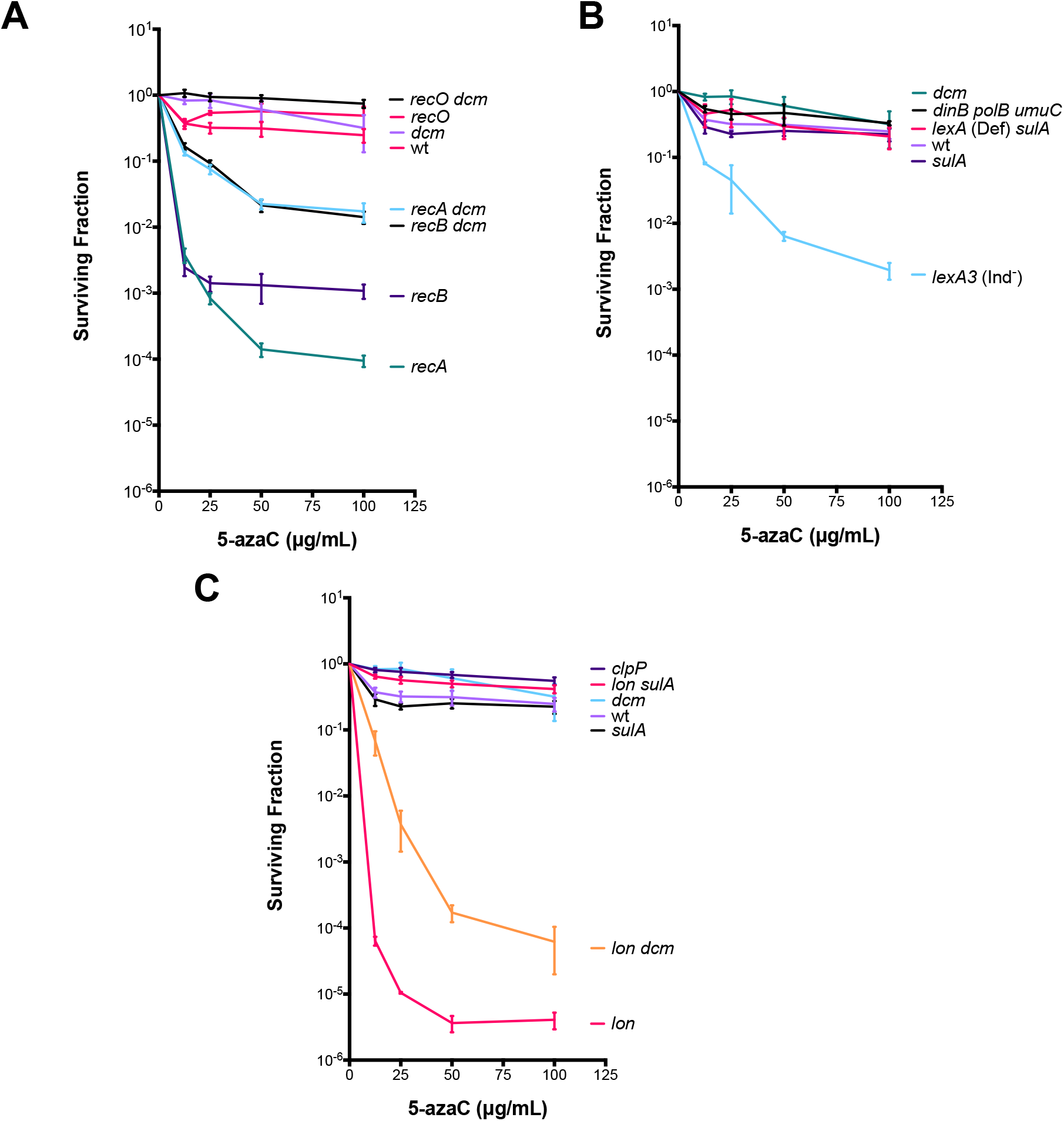
Fractional survival of *E. coli* genetic variants at indicated doses of 5-azaC for 2 hours. Data are average of at least 3 experiments; error bars represent standard deviation. A. Recombination mutants B. SOS response mutants C. Protease mutants.

**Figure 3.**
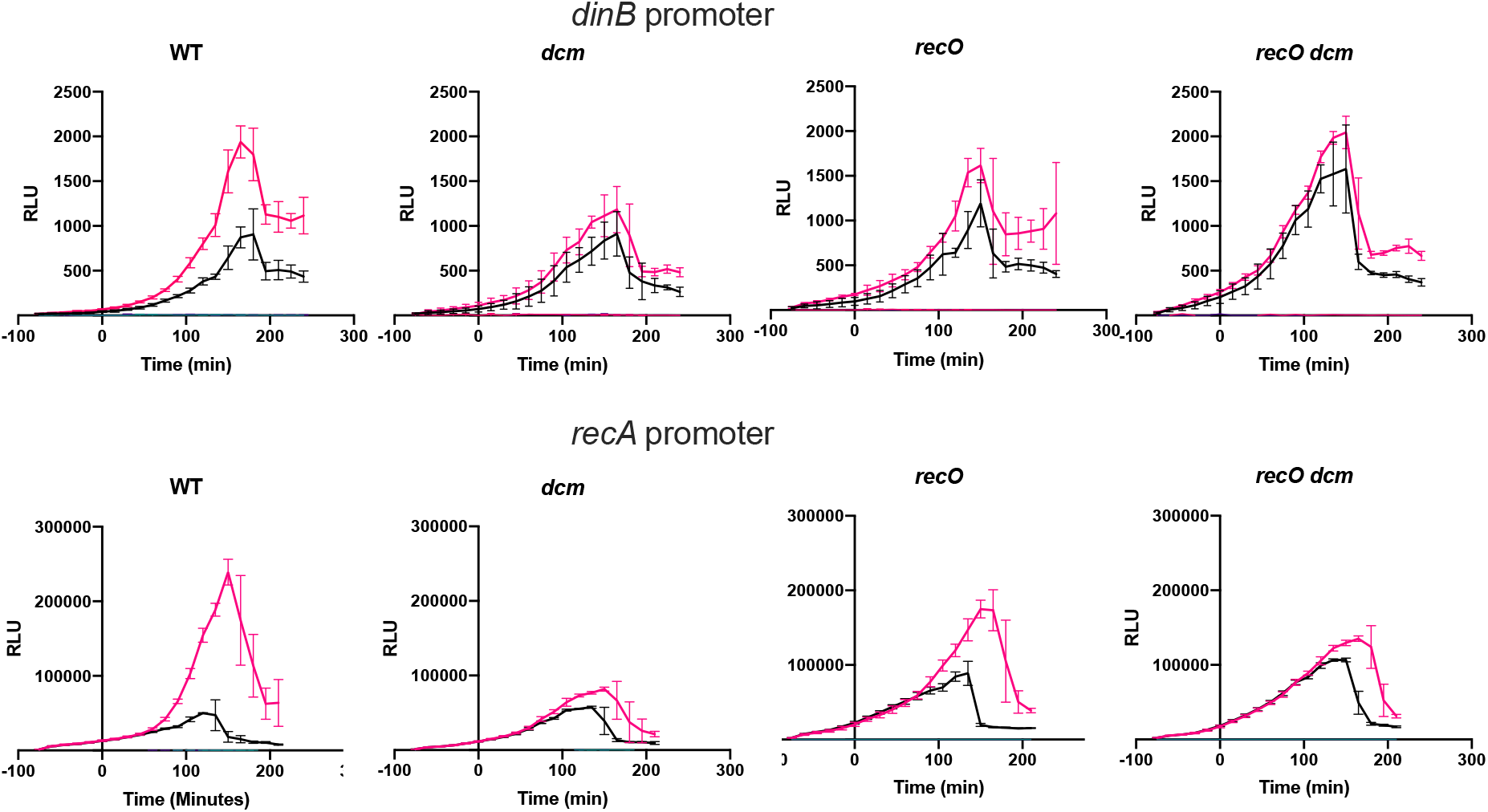
Induction of SOS promoters, *dinB* and *recA*, fused to luciferase expression reporters. Red treated with 50 μg/ml 5-azaC; black untreated. 5-azaC was added at time 0 and the culture was followed for 300 minutes post-treatment. RLU, relative luciferase units is the luminescence divided by the OD_600_ of the culture.

RecA is required for homologous recombination in *E. coli*, and it participates in two genetically distinct pathways. One of these, involving the RecBCD nuclease, is specific for recombination arising from DSBs, the other, dependent on RecFOR, is specific for recombinational repair of single-strand gaps (reviewed in (Persky and Lovett 2008)). These pathways are defined by the functions that facilitate the loading of RecA, which RecBCD performs on newly resected DNA and RecFOR on ssDNA gaps, the latter displacing bound single-strand DNA binding protein (SSB). The RecA filament bound to ssDNA not only directly facilitates homology search and DNA strand exchange, but is also the inducing signal for the SOS response, a transcriptional response in *E. coli* to DNA damage (Simmons et al. 2008; Lovett 2010). We show here that RecO mutants exhibit no sensitivity to 5-azaC (Fig. 2A) unlike mutants in RecA or RecBCD.

One difficulty in assessing the role of the RecFOR pathway is that, in its absence, gaps become converted to DSBs and are repaired efficiently by the RecBCD pathway. By survival assays to genotoxins, there may be no evidence for RecFOR participation. However, genetic measurements of homologous recombination and induction of the SOS response may show that the RecFOR pathway is indeed called into play. For example, the replication chain terminator, azidothymidine (AZT), blocks replication and induces the SOS response through the RecFOR pathway, although *recFOR* mutants show no sensitivity to AZT (Cooper and Lovett 2010).

To ascertain the participation of the RecFOR pathway in processing DPCs, we measured SOS induction in *E. coli* after 5-azaC treatment, using a luciferase transcriptional reporter fused to the *dinB* and *recA* promoters. Both of these genes are regulated as part of the SOS regulon, although *recA* is more highly expressed than *dinB*. In these assays, cells are continuously exposed to sublethal concentrations of 5-azaC, beginning in early log phase and monitored through the growth of the culture into stationary phase. Luminescence of the culture is normalized to OD6_00_ of the culture, measured at 15 minute intervals. 5-azaC does indeed induce expression of both *dinB* and *recA* promoters, reaching a maximum about 2 hrs after introduction of the drug. This induction was dependent on *dcm*, confirming that DPCs are responsible for the induction. For both induced and uninduced expression, we saw a growth-dependent increase in expression, peaking at late exponential phase. We do not understand the basis of the effect. This increase was not seen in control cultures with the luciferase operon, lacking the promoter regions, so this signal is dependent on the *recA* or *dinB* promoters. We saw a modest reduction by *recO*, although the effect is obscured by the elevation of expression in *recO* untreated cultures, possibly due to the accumulation of spontaneous DNA lesions in this genetic background.

### 3.2 Does 5-azaC induce recombination, as well as the SOS response?

We measured two types of recombination with genetic assays: one selects for crossovers between 411 bp of homology on low copy plasmids (Lovett et al. 2002) and the other detects nonreciprocal gene conversion between an internally deleted *lacZ* gene and an ectopic internal ‘*lacZ*’ gene segment at a position across the chromosome, near *oriC* (see Materials and Methods).

Crossovers detected in the plasmid assay are primarily dependent on the RecFOR pathway (Lovett et al. 2002); because these plasmids lack Chi sites that protect from degradation by RecBCD (Anderson and Kowalczykowski 1997); there is no detectable contribution of the RecBCD pathway. Using cultures that were split and treated for 2 hr with 5-azaC (Fig. 4), we saw a modest 2-to 3-fold elevation of crossing-over with 5-azaC relative to untreated cultures. This increase was dependent on *recO*. 5-azaC induced recombination was dependent on *dcm* as well, with *dcm* mean fold-induction values of 0.89 ± 0.18, 0.60 ± 0.21, 0.50± 0.39 at 25, 50 and 75 μg/ml 5-azaC, respectively (n=4).

**Figure 4.**
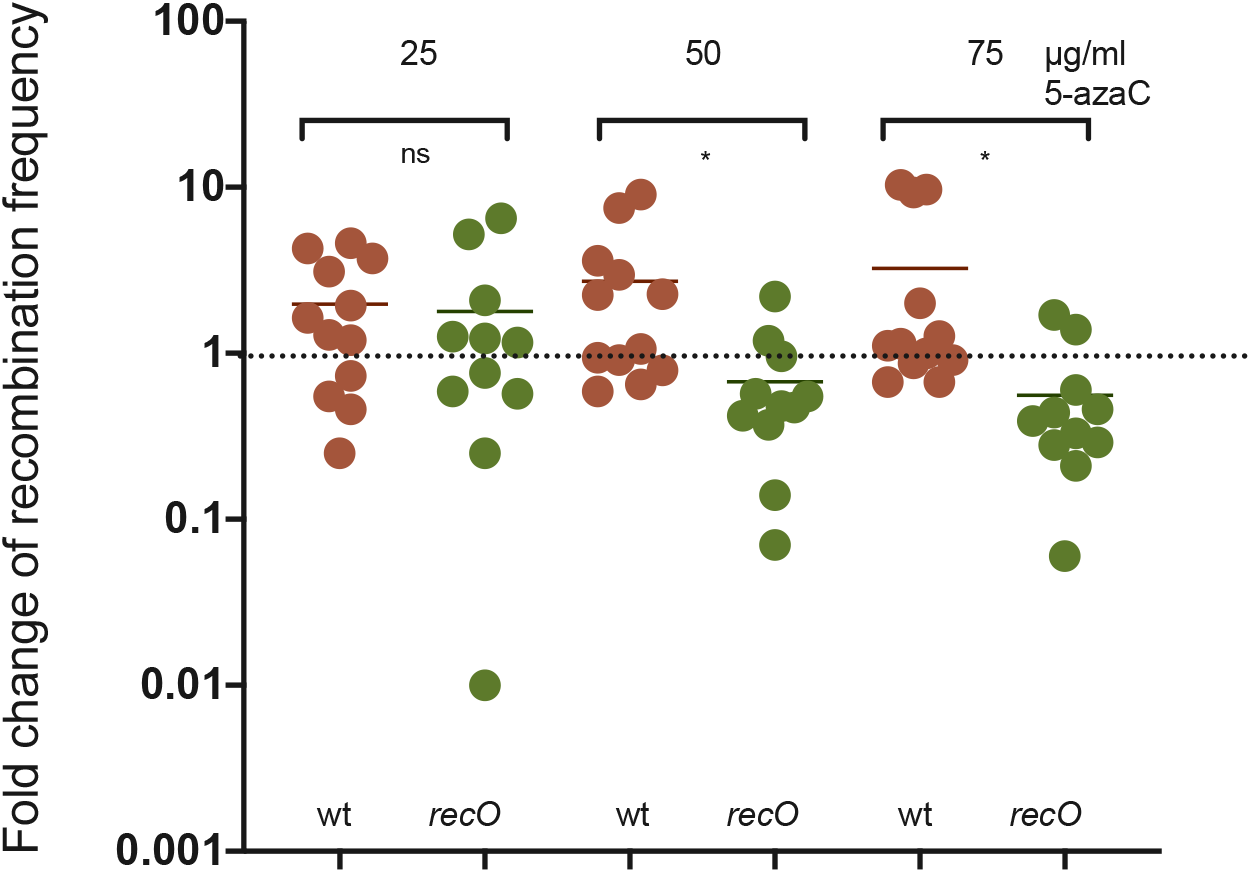
Crossover recombination induced by 5-azaC, 2 hr treatment at indicated doses, in split cultures of wt or *recO* mutant strains. Bars represent mean values of the replicates.

Gene conversion between chromosomal *lacZ* alleles was more strongly stimulated by 5-azaC, with an over 30-fold increase at 50 μg/ml. (Figure 5). This was reduced 2-fold by *recO*, although there was considerable variation among replicates. We saw no evidence for Dcm-dependence of this recombination, so there must be a recombinogenic lesion induced by 5-azaC other than Dcm-DPCs. The residual recombination in *recO* mutants is presumedly dependent on the alternate RecBCD recombination pathway (Persky and Lovett 2008), but we were unable to test *recB* and *recB recO* mutants in this assay, due to their strong sensitivity to 5-azaC.

**Figure 5.**
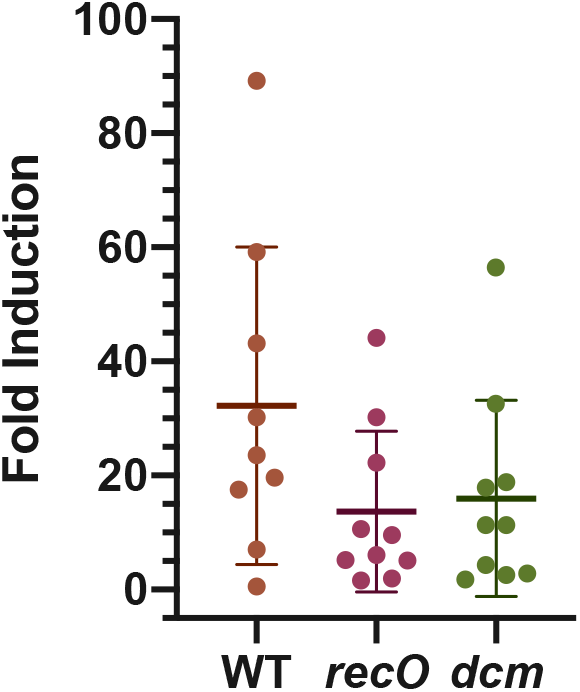
Gene conversion recombination induced by 50 μg/ml 5-azaC, 2 hr treatment at indicated doses, in split cultures of wt or recO mutant strains. Bars represent mean values of the replicates and standard deviations.

### 3.3 Template-switching at quasipalindromes: effect of recombination functions

We had previously documented a strong stimulatory effect of 5-azaC on mutagenesis detect by template-switching in a quasipalindromic sequence. In this study, we expanded this analysis to include additional mutants and doses of 5-azaC. We examined, in parallel, the effect of this set of homologous recombination factors on QPM using leading and strand strand reporter strains, measuring *lacZ* reversion frequencies with and without treatment with 5-azaC at two doses. For each genetic background, an isogenic *dcm* derivative was also assayed. There are 8 potential Dcm methylation sites within *lacZ*. As we had seen previously (Laranjo et al. 2018; Klaric et al. 2020), we observed stimulation of QPM with increasing 5-azaC dose, with a stronger effect with the leading strand template-switch reporter QP5. (Fig. 6) Most, but possibly not quite all, of this induction is lost in *dcm* mutants. Mutants in *recA* showed no decrement in mutagenicity; if anything, they showed a slightly higher constitutive and 5-azaC-induced frequencies, at both doses, for both reporters. Mutants in *recA* had a strong effect of 5-azaC on the lagging strand reporter QP6 where it elevated QPM more than an order of magnitude at the higher dose. Again, as for wt, 5-azaC-induced QPM in *recA* was largely absent in *dcm* double mutants. Mutants in *recB* showed a slightly higher constitutive rate of QPM, but 5-azaC induction could not be tested at the higher dose because of poor survival. Mutants in *recO* showed a higher constitutive frequency of QPM in untreated cells but looked virtually indistinguishable from wt after 5-azaC treatments. These results suggest that homologous recombination and/or SOS induction promotes higher survival to 5-azaC, with RecA playing a antimutagenic role for QPM. Elevated constitutive QPM rates for *recB* and *recO* mutants would be consistent with the idea that both RecBCD and RecFOR pathways process spontaneous lesions, with a stronger effect of the latter pathway.

**Fig. 6.**
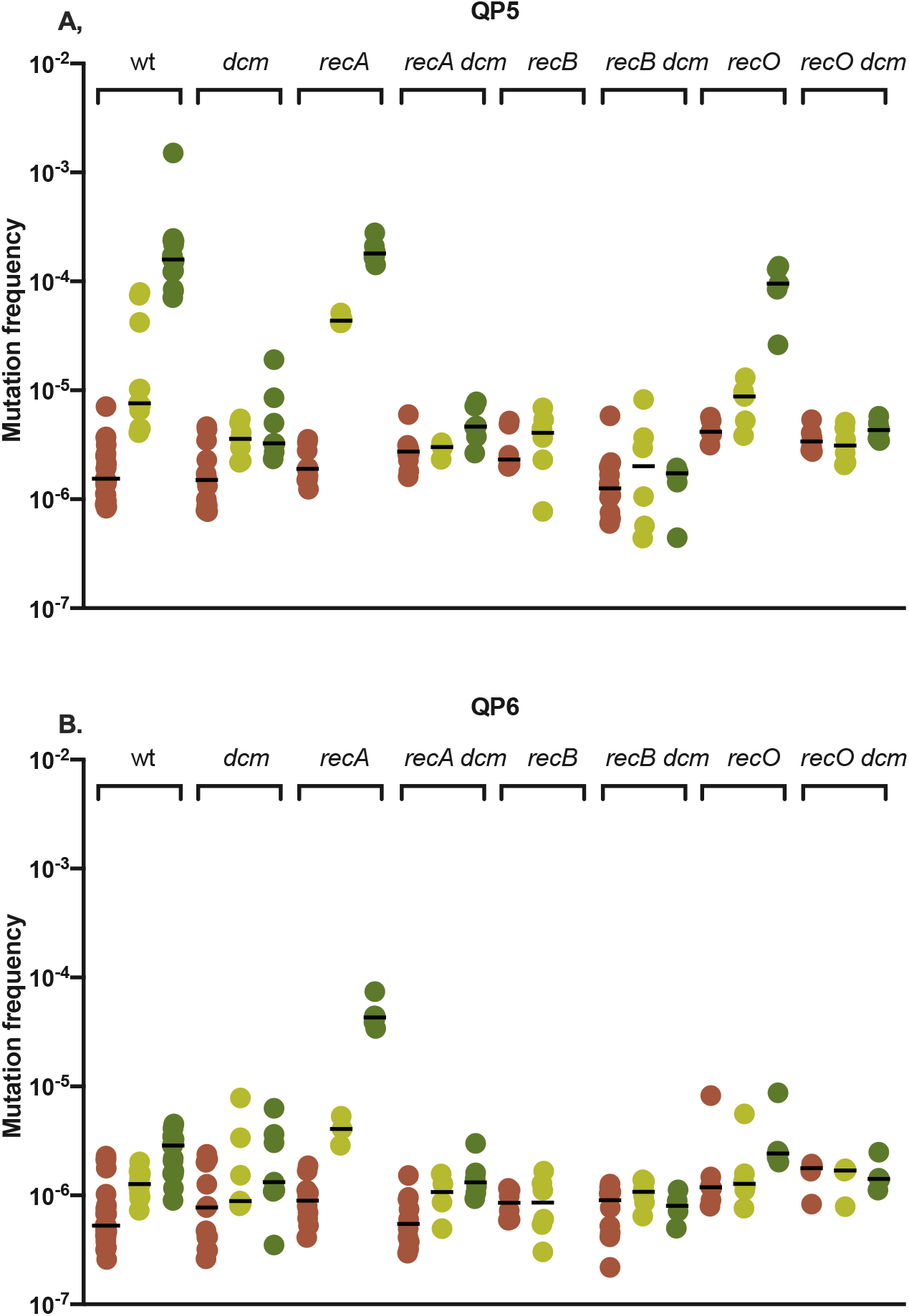
Mutation frequencies with and without 5-azaC treatment in recombination mutants, genotype shown above bracket. Shown are *dcm*^+^ and *dcm* derivatives of wt, *recA*, *recB* and *recO* mutants. QP5 is a leading strand mutational reporter; QP6 a lagging strand reporter. The log scale of the the Y-axis is the same in the two graphs, to allow easier visual comparison. No treatment frequencies in brown, 1 μg/ml 5-azaC in light green, 12.5 μg/ml in dark green. Black bar indicates the median value of the replicates.

### 3.4 Template-switching at QPs: effect of SOS response and translesion DNA polymerases

The SOS response of *E. coli* consists of a number of functions coordinately induced by DNA damage or replication inhibition (Simmons et al. 2008). The genes for these functions are transcriptionally repressed by LexA, but which are induced by LexA self-cleavage promoted by the accumulation of RecA/ssDNA filaments. By measuring QPM in *lexA3* strains, in which the SOS response cannot be induced, we had previously shown that the SOS response is anti-mutagenic for QPM, both constitutively and that induced by DPCs ((Laranjo et al. 2018) and Figure 8.) Mutants in *lexA3* are also sensitive to killing by 5-azaC (Figure 2B). Several of the factors induced by the SOS response are the translesion DNA polymerases, Pol II, Pol IV and Pol V (encoded by *polB, dinB* and *umuCD* genes, respectively). To determine if these were responsible for the antimutagenic effect of the SOS response, we mutated all three and measured spontaneous QPM rates in a wild-type genetic background and in strains lacking one or both of the 3’ to 5’ exonucleases Exo I and Exo VII.

**Figure 7.**
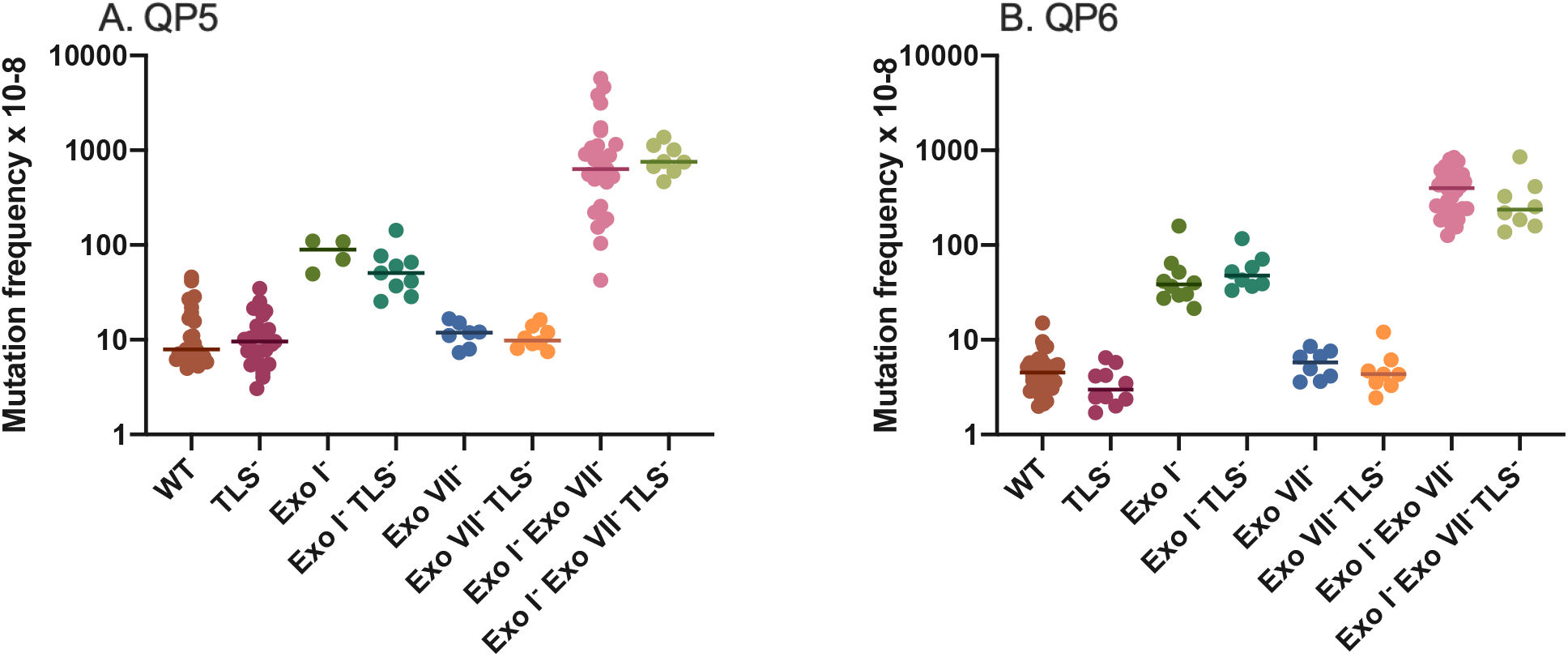
QPM frequencies in strains lacking the TLS DNA polymerases (*polB dinB umuC*) with and without the DNA exonucleases Exo I and/or Exo VII. Bars indicate the median value of the replicates.

**Fig. 8.**
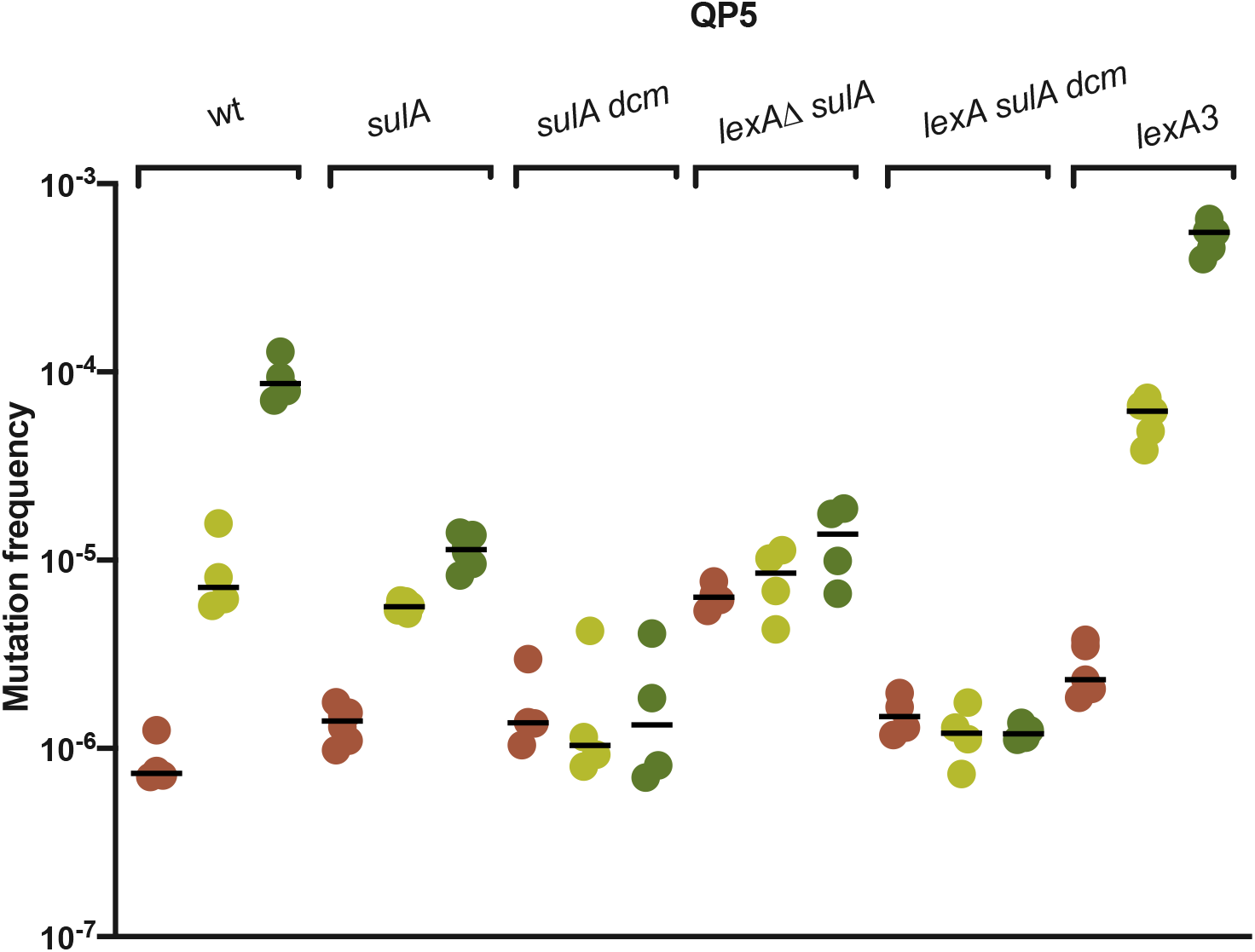
Mutation frequencies with and without 5-azaC treatment in *lexA* mutants, genotype shown above bracket. Shown are *dcm*^+^ and *dcm* derivatives of *sulA* and *lexA*Δ *sulA* compared to WT and *lexA3* mutants assayed in parallel. QP5 is a leading strand mutational reporter. No treatment frequencies in brown, 1 μg/ml 5-azaC in light green, 12.5 μg/ml in dark green. Black bar indicates the median value of the replicates.

These exonucleases abort QPM, with ExoI playing the larger role (Laranjo et al. 2017); hence QPM rates are much higher in their absence and potentially more sensitive to TLS polymerase effects. We saw no effect in any background of TLS polymerases on mutation frequency, for either the leading or lagging strand QPM reporters (Figure 7). The loss of all three TLS polymerases also did not reduce survival to 5-azaC (Figure 2B.) Therefore, induction of TLS is not responsible for the antimutagenic aspect of the SOS response.

We examined mutants in which the SOS response was constitutively induced, lacking the LexA repressor. Because the cell division inhibitor SulA is induced during the SOS response, *lexA*Δ strains must carry an additional knockout of *sulA* to remain viable. With the QP5 reporter, *lexA sulA* null strains showed diminished 5-azaC-induced QPM relative to wt strains (Figure 8). However, the *sulA* mutation by itself appeared to depress mutagenesis at the higher 5-azaC concentration (Fig.8). One explanation for this effect is that potential QP mutants require SulA-dependent cell division arrest to recover and form a colony. The 5-azaC-induced mutagenesis was entirely dependent on *dcm* (Fig. 8), showing that it is promoted by the DPCs formed by Dcm methyltransferase in the presence of 5-azaC.

The most striking effect of *lexA*Δ, however, was a more than 10-fold elevation of constitutive rates of QPM (in the absence of 5-azaC) relative to wt and *sulA* strains (Fig. 8). This suggests that the SOS response also can *promote* mutagenesis in some instances.

We wondered whether the unnatural, full-on induction of TLS DNA polymerases was responsible for this effect. We examined strains overexpressing one of the TLS DNA polymerases from an arabinose-regulated pBAD plasmid, with a copy number approximately 20 per cell, in an otherwise wt strain. In the absence of induction, we saw no effect of the TLS polymerases. After 2 hours of induction in arabinose medium, there was a strong mutagenic effect of DNA polymerase IV (DinB) for mutations detected by both QP5 and QP6, 34- and 13-fold, respectively. (Figure 9). Pol IV is known to be prone to −1 G frameshift mutations (Kim et al. 1997) and in a *lac* reporter specific for this mutation, we saw a very large increase in mutation frequency by Pol IV when induced. Induction of Pol II caused about a 3-fold increase in QPM on the leading strand (QP5), and a 2-fold effect on the lagging strand (QP6). Pol V also was modestly mutagenic. This suggests that the TLS polymerases are mutagenic for QPM when their abundance is unbalanced, particularly Pol IV, a potential explanation for the hypermutable constitutive phenotype of *lexA*Δ *sulA* strains.

**Figure 9.**
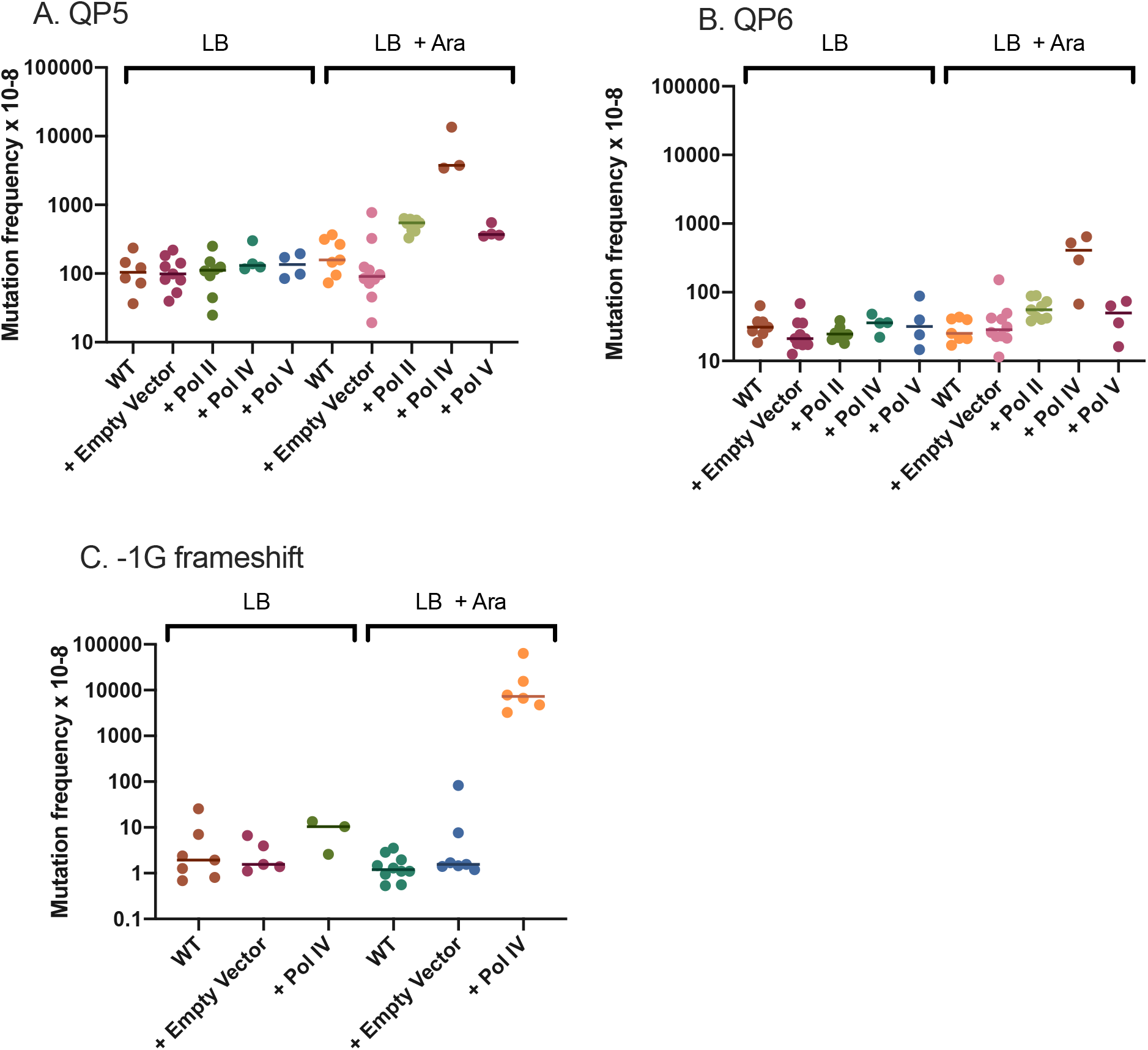
Induced expression of TLS DNA polymerase and effects on mutation frequencies with the indicated *lacZ* mutational reporter strains. Bars indicate the median value of the replicates.

### 3.5 Template-switching at QPs: effects of proteases

Because DPCs may be removed by proteolysis (Vaz et al. 2017), we examined two major intracellular, energy-dependent proteases of *E. coli* for effects on 5-azaC induced QPM. Mutants in the Lon protease were found to be extremely sensitive to killing by 5-azaC (Figure 2C) and this sensitivity was abolished completely by a secondary mutation in the cell division inhibitor, *sulA*. During recovery from the SOS response, Lon protease destroys SulA, allowing cells to divide again. In the absence of Lon, SOS induction is lethal because cells never recover the ability to divide (Schoemaker et al. 1984). Complete suppression of *lon* by *sulA* (Figure 2C) suggests that the failure to turn off the SOS response to resume cell division is predominantly responsible for 5azaC-induced killing in this background. A secondary mutation in Dcm conferred only partial resistance to 5-azaC in *lon* mutants. Therefore, some lethal SOS induction derives from Dcm DPCs formed with 5-azaC, but there must be a second SOS-inducing lesion that is Dcm-independent. Mutants in ClpP, a component of both ClpAP and ClpXP protease complexes (Gottesman 1996), did not show any sensitivity to 5-azaC (Figure 2C).

With QPM reporter QP5, mutants in *lon* showed hypermutability, 20-fold relative to wt, after treatment with the lower dose of 5-azaC (the higher dose could not be tested because of killing) (Figure 10). This elevated mutagenesis was entirely dependent on *dcm* (Figure 10). Mutants in *clpP* exhibited normal levels of Dcm-dependent QPM as assayed with reporter QP5.

**Fig. 10.**
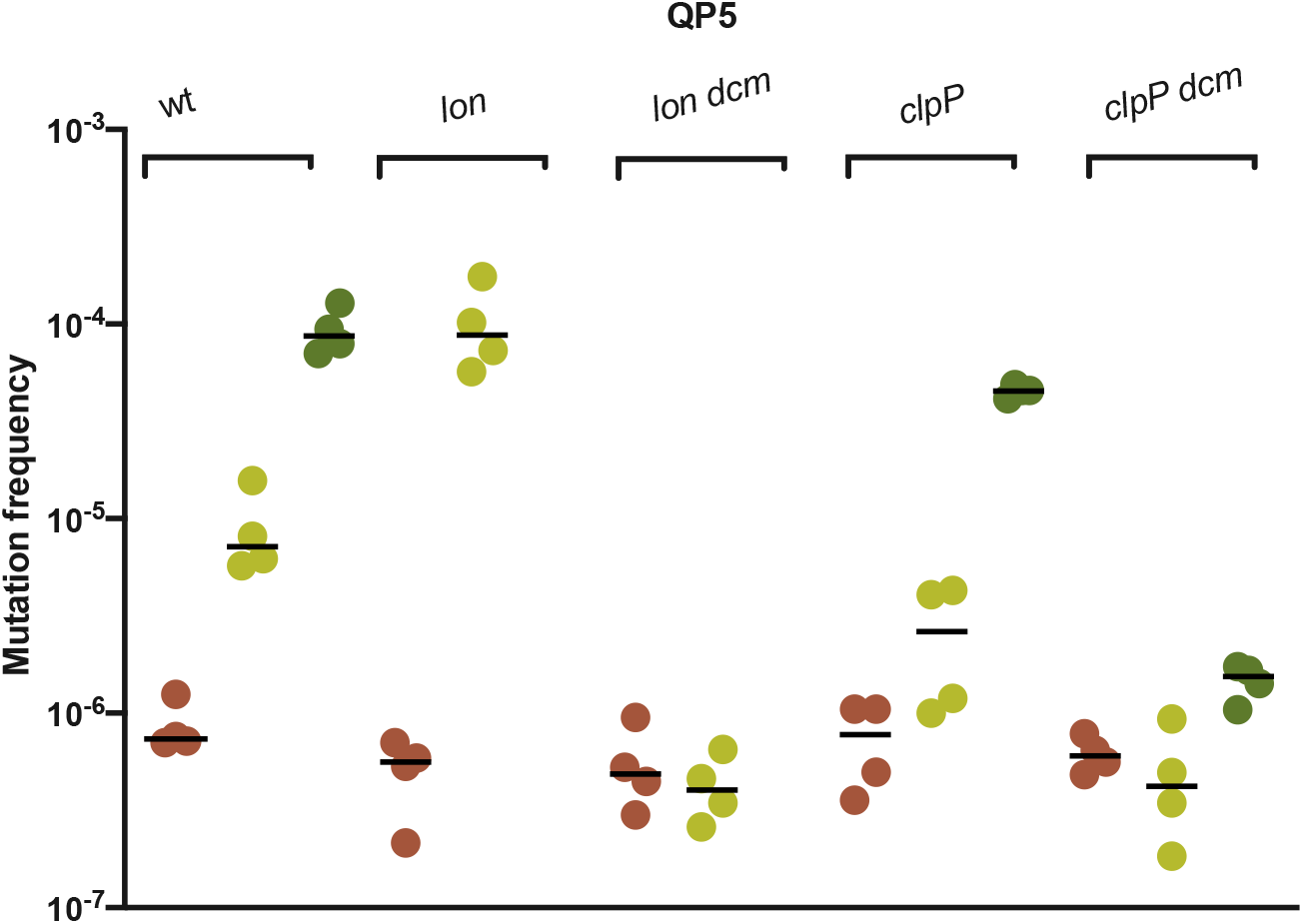
Mutation frequencies with and without 5-azaC treatment in *lon and clpP* mutants, genotype shown above bracket. Shown are *dcm*^+^ and *dcm* derivatives of *lon* and *clpP* mutants, compared to wt assayed in parallel. QP5 is a leading strand mutational reporter. No treatment frequencies in brown, 1 μg/ml 5-azaC in light green, 12.5 μg/ml in dark green. Black bar indicates the median value of the replicates.

### 3.6 Template-switch mediated deletions

Deletions and expansion of tandem direct repeats occurs by a template-switch mechanism, independent of homologous recombination proteins including *recA* (reviewed in (Lovett 2017)). With 2 different assays for deletion formation (Lovett et al. 1994; Seier et al. 2011), we tested the effects of DPC-agents 5-azaC and formaldehyde. 5-azaC showed a dosedependent increase in the deletion frequency of 11 bp tandem repeats within the chromosomal *lac* locus, with a 4-fold enhancement of deletion at the highest dose of 100 μg/ml (Figure 11a). Note that with a hundred-fold lower dose of 1 μg/ml 5-azaC, we saw a 5-fold enhancement of QPM *lac* mutagenesis using the QP5 reporter (Fig. 3A), so template-switching that gives rise to short deletions is not as strongly affected by 5-azaC as are QP-templated mutations. The 5-azaC effect was reduced to 2-fold in the *dcm* mutant (Fig. 11b), although the distributions considerably overlap. This indicates that although some of the deletionogenic effect of 5-azaC may occur through Dcm DPCs, some 5-azaC-induced deletions are independent of Dcm. Formaldehyde had a much stronger effect on deletion in this assay, with an enhancement of over 20-fold in the standard 2-hour treatment (Figure 11a). In a plasmid assay for deletion of 101 bp repeats within the *tetA* gene, conferring tetracycline resistance, we saw that 2 hr of 5-azaC treatment enhanced deletion frequencies 2-fold, which was reduced to 1.2 fold in the *dcm* mutant (Figure 11c). Similar treatment of wt type cells with formaldehyde enhanced deletion in this assay 3-fold and it was not appreciably affected by *dcm*, as expected, since the DPCs induced by FA are nonspecific.

**Fig. 11.**
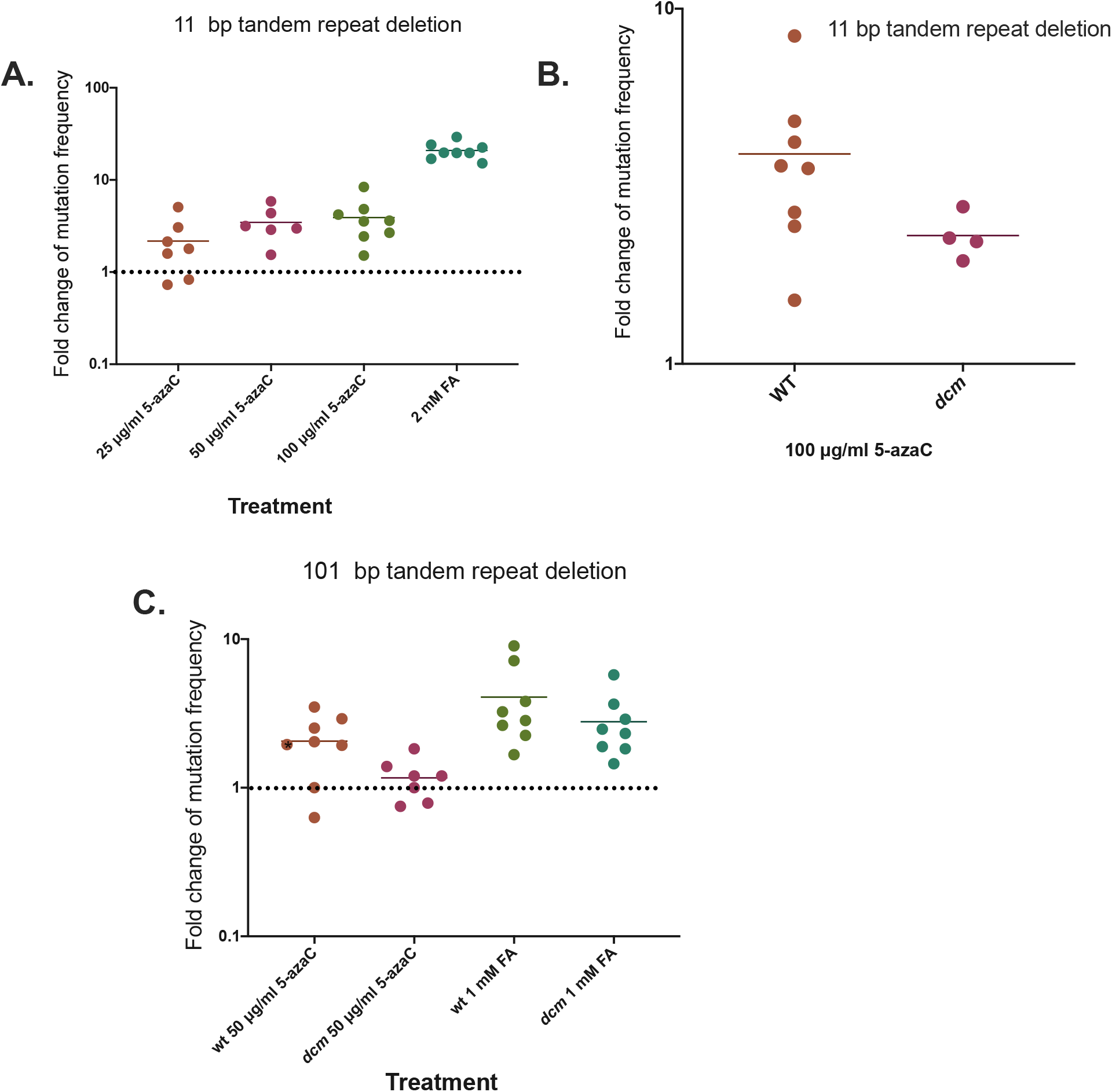
Tandem repeat deletion induced by 5-azaC and FA. Cultures were split and treated for 2 hr with 5-azaC or FA. The mutation frequency in the treated culture relative to that in the untreated is plotted: a value of 1 (indicated by dashed line in A and C) means there is no effect. Bars indicate mean values. A. Deletion of 11 bp repeats in chromosomal *lacZ*. B. Effect of *dcm* on 11 bp *lacZ* deletion. C. Deletion of 101 bp repeats on plasmid-encoded *tetA* and effects of *dcm*.

## 4. DISCUSSION

We used 5-azacytidine to investigate the genetic consequences of DPCs formed by cytosine methytransferase, Dcm, in *E. coli*. Using a variety of genetic assays, we measured 5-azaC effects on the DNA damage response, recombination and genetic mutations produced by template switching, at inverted or direct DNA repeats.

### 4.1 The DNA damage response

By transcriptional reporter assays for *dinB* and *recA* promoters, we show that 5-azacytidine induces the SOS response, a transcriptional response triggered by the formation of RecA filaments on single-strand DNA in vivo. This induction was dependent on Dcm, implicating the Dcm protein-DNA crosslink as the inducing lesion. We saw a modest reduction in a RecO mutant, implicating the RecFOR pathway for SOS induction; however, at least for the RecA promoter, there was detectable 5-azaC induction independent of RecO. We suspect that the RecBCD pathway also induces SOS in the absence of RecO, but this dependence could not be tested due to poor survival of *recB* strains to 5-azaC.

### 4.2 Homologous recombination

A 2-hour treatment with 5-azaC induced homologous recombination about 30-fold, as detected by a gene conversion assay between an internally-deleted *lacZ* gene and a 500 bp homologous *lacZ* fragment integrated across the chromosome, at *att*Tn*7*. This increase in recombination was also seen in a *dcm* mutant, suggesting that lesions other than Dcm-DNA crosslinks can initiate recombination. There was a small, 2-fold, reduction in 5-azaC-induced recombination in a *recO* mutant. A plasmid-based assay for crossover recombination, showed a more modest 2-3 fold stimulation by a 2-hr treatment with 5-azaC, which was reduced by *recO* and *dcm*. Therefore, we found evidence for homologous recombination induced by 5-azaC, some associated with DPCs and some not.

### 4.3 Mutability at QP sites/ effects of recombination

In the absence of RecA, 5-azaC mutagenicity was enhanced at quasipalindromic (QP) sites, dependent on the Dcm methyltransferase. The elevation by *recA* was particularly striking for the QP6 reporter for lagging-strand template-switching, where loss of *recA* elevated mutation frequencies, relative to wt, 8- and 68-fold at 1 and 12.5 μg/ml 5-azaC, respectively. This elevation of lagging strand QPM was not due to lack of the SOS response, since *lexA3* mutant strains, likewise unable to mount the SOS response, showed no or 3-fold elevation of QP6 rates at 1 and 12.5 μg/ml 5-azaC, respectively, relative to wt type strains. Therefore, the capacity for homologous recombination, negated by *recA* but not by *lexA3*, helps avoid QPM by 5-azaC but apparently only, or predominantly, on the lagging strand.

### 4.4 Mutability at QP sites/ effects of the SOS response and TLS DNA polymerases

The SOS response does have an antimutagenic effect on 5-azaC -induced leading strand QPM, with 4-6-fold enhancement by *lexA3* and 8-fold by *recA*, relative to wt. Our prior work with the natural QP mutational hotspot in the *thyA* gene had indicated that the TLS polymerases, under SOS control, had an antimutagenic effect (Dutra and Lovett 2006), a potential explanation for the antimutagenic nature of the SOS response. However, in this work, using QP sites within the *lacZ* gene, we can find no evidence for such an effect. We saw no effect on mutation rates by introduction of *polB dinB umuC* knockouts, even when ExoI and ExoVII were absent. Rather, these SOS-induced polymerases, when overexpressed, appeared to be mutagenic for QPM. When expressed from the arabinose pBAD promoter on a medium copy number plasmid, induction of any of the three polymerases elevates QPM, with the strongest effect by *dinB*. Because we do not know the extent of the induced expression, this is difficult to interpret, although it does indicate that unbalanced polymerase expression has the potential to cause QPM. This is also consistent with our finding that *lexA*Δ strains show an elevated spontaneous level of QPM. Although Pol IV (DinB) is reported to have a greater mutagenic effect on the lagging strand (Fijalkowska et al. 2012), we see increases in QPM with both leading and lagging strand reporters, although this could be influenced by over-expression.

Surprisingly, a *sulA* mutation, which prevents cell division arrest after DNA damage, reduced the mutagenicity of 5-azaC, as detected on the leading strand QP5 reporter, from 3-to 5-fold. We know of no precedent for *sulA* effects on mutagenesis, with the exception of a very modest (<2x) effect on stress-induced mutation (Al Mamun et al. 2012). Except for *dcm* mutants, this is the only strain to exhibit lower frequencies of mutation than wt after 5-azaC treatment. We can explain the lower yield of mutants in the *sulA* strain by proposing that potential mutants fail to survive in the absence of cell division arrest. and so are lost in this assay. The loss of *sulA* did not, however, influence survival of the bulk cell population to 5-azaC, so the vulnerable cells in this case must be a subpopulation.

### 4.5 Mutability at QP sites/ effects of proteases

The Lon protease plays a potential role in the avoidance of 5-azaC induced mutations at quasipalindromes; *lon* mutants were hypermutable by over 50-fold at the lower 5-azaC dose. We documented this elevation using the leading strand QPM reporter, but saw a similar enhancement on with the lagging strand reporter (Klaric, unpublished results). The persistence of a DPC in the absence of Lon may promote template-switching, whereas prompt removal of the DPC by Lon proteolysis leads to a nonmutagenic outcome. Mutants in *lon* were extremely sensitive to killing by 5-azaC, but this appears to be entirely due to the failure to recover after induction of the SOS response; this sensitivity was suppressed completely by an additional mutation in *sulA*. We saw no effect by ClpP.

### 4.6 5-azaC: not just DPCs?

Our data suggest that although Dcm-dependent DPCs largely contribute to genetic effects of 5-aza, there is some Dcm-independent lesion(s) as well. This is particularly apparent in the survival curves of strains sensitive to killing by 5-azaC (*recA*, *recB*, and *lon* mutants), where *dcm* improves survival but does not restore it to wildtype levels. In addition, homologous recombination assayed by gene conversion between *lacZ* alleles is induced strongly by 5-azaC but is largely *dcm*-independent; 5-azaC-induced deletion of 11 bp in *lacZ* is partially *dcm*-independent. Whatever the Dcm-independent lesion is, it must induce the SOS response since *sulA* entirely suppresses 5-azaC killing of *lon* mutants, whereas *dcm* suppresses it only partially. Dcm-independent killing of *recA* and *recB* mutants by 5-azaC suggests that the lesion must also lead to DSBs in DNA. 5-azaC can break down into guanylurea (Figure 1), which in DNA mispairs with cytosine, leading to C to G transversion mutations (Jackson-Grusby et al. 1997). In mammalian cells, this C to G transversion correlates with sites of C-methylation, leading to speculation that the CMeT induces the cytosine ring opening that converts it to guanylurea (Jackson-Grusby et al. 1997). However, more recent work suggest that guanylurea deoxribonucleotides can be directly incorporated into DNA during replication (Lamparska et al. 2012). Using a specific reporter for C to G transversion mutations (Cupples and Miller 1989; Seier et al. 2011), we confirmed that 5-azaC does indeed have a strong, 500-fold mutagenic effect on C to G transversion (confirming (Cupples and Miller 1989)), and occurs entirely independent of *dcm* (Table S1 and (Doiron et al. 1999)). Therefore, guaunylurea lesions may indeed be a potential contributor to *dcm*-independent effects of 5-azaC, where it may act like a abasic lesion to block DNA synthesis or elicit processing that leads to mutagenesis, template switching or recombination. It is also possible that there are additional unknown CMeTs encoded in the *E. coli* genome that are responsible for the *dcm*-independent effects.

### 4.7 Differential effects on template-switching

Although a template-switch mechanism has been proposed for both QPM at inverted repeats and deletions/expansion at direct repeats (reviewed in (Lovett 2017), we saw a much stronger effect on the former than the latter. We assayed two types of deletions that arise between short tandem repeated sequences, 11 bp in the chromosomal *lacZ* gene, and 101 bp in the *tetA* gene on a ColE1 plasmid and both were stimulated modestly, 2-4 fold after 2 -hr treatment with 5-azaC. In contrast, comparable treatment yielded stimulation of QPM over 2 orders of magnitude. We do not know the basis for this differential effect but further work may clarify the differences between these template-switching events.

## 5. Acknowledgments

We thank Hirotada Mori and the National Genetics Institute of Japan for the collection of E. coli knockout mutants. Funding: This work was supported by the National Institutes of Health, grants R01 GM51753 and P01 GM105473 to STL, T32 GM007122 to JAK and National Science Foundation REU grant DBI-1359172 to ELP.

---

**Table S1.** Frequency of C to G transversions with and without 5-azaC, in wild-type and dcm mutants, n= 4

|  | **Mutation frequency x 10<sup>-8</sup>** |  |
| --- | --- | --- |
|  | **wt** | **dcm** |
| **untreated** | 0.052 | <0.05 |
| **100 µg/ml 5-azaC** | 25.3 | 25.6 |

## Footnotes

1 Abbreviations: DPCs: DNA protein crosslinks CMeT: cytosine methyltransferase 5-azaC: 5-azacytidine DSBs: double strand breaks ssDNA: single-strand DNA QP: quasipalindrome QPM: quasipalindrome-associated mutation

## Notes

### Competing Interest Statement

The authors have declared no competing interest.

